# Dynamic post-transcriptional regulation during embryonic stem cell differentiation

**DOI:** 10.1101/123497

**Authors:** Patrick R. van den Berg, Bogdan Budnik, Nikolai Slavov, Stefan Semrau

## Abstract

During *in vitro* differentiation, pluripotent stem cells undergo extensive remodeling of their gene expression profile. While studied extensively at the transcriptome level, much less is known about protein dynamics. Here, we measured mRNA and protein levels of 7459 genes during differentiation of embryonic stem cells (ESCs). This comprehensive data set revealed pervasive discordance between mRNA and protein. The high temporal resolution of the data made it possible to determine protein turnover rates genome-wide by fitting a kinetic model. This model further enabled us to systematically identify dynamic post-transcriptional regulation. Moreover, we linked different modes of regulation to the function of specific gene sets. Finally, we showed that the kinetic model can be applied to singlecell transcriptomics data to predict protein levels in differentiated cell types. In conclusion, our comprehensive data set, easily accessible through a web application, is a valuable resource for the discovery of post-transcriptional regulation in ESC differentiation.

## Introduction

Much of the medical potential of pluripotent stem cells is due to their ability to differentiate *in vitro* into all tissue types of the adult body (Soldner and Jaenisch, 2012). While tremendous progress has been made in guiding cells through successive lineage decisions, the gene regulatory mechanisms underlying these decisions remain largely unknown. This gap in knowledge hampers the streamlining and acceleration of differentiation protocols. A large body of work has focused on transcriptional regulation, charting transcriptome changes during differentiation, most recently down to the single-cell level (Klein et al., 2015; Loh et al., 2016; Semrau et al., 2016) These studies assumed implicitly that mRNA levels are a good proxy for protein levels. Mounting evidence suggests that this is not a good assumption for mammalian systems, where mRNA and protein levels were found to correlate only moderately (Lu et al., 2009) (Kristensen et al., 2013; Peshkin et al., 2015; Schwanhäusser et al., 2011). Where the discordance between protein and mRNA expression originates and what the biological function might be are long-standing and controversially discussed issues (Liu et al., 2016; Vogel and Marcotte, 2012). Here we study the relationship between mRNA and protein expression in the context of *in vitro* differentiation, a highly dynamic process in which gene regulation at the protein level likely plays an important role (Sampath et al., 2008).

## Results

### Measurement of transcriptome and proteome dynamics during retinoic acid driven differentiation

We used retinoic acid (RA) differentiation of mESCs as a generic model for *in vitro* differentiation. Previously, we characterized this differentiation assay in detail at the transcriptional level by single-cell RNA-seq (Semrau et al., 2016). In particular, we have shown that within 96 h of RA exposure, mESCs bifurcate into an extraembryonic endoderm-like and an ectoderm-like cell type (XEN and ECT respectively). Here we collected samples of the mixed population during an RA differentiation time course as well as the two final, FACS-purified differentiated cell types at 96 h (Fig. 1a). For each time point or cell type we quantified poly(A) RNA by RNA-seq and protein expression by tandem mass tag (TMT) labeling followed by tandem mass spectrometry (MS/MS). In total, we obtained both RNA and protein expression for 7459 genes (Supplementary Fig. 1a). Protein levels were quantified with low technical error (Supplementary Fig. 1a) and high reproducibility between protein fold changes measured in biological replicates (Pearson’s r = 0.92, Supplementary Fig. 1b). Moreover, the XEN-like cells measured here were similar to embryo derived XEN cells in their proteome (Mulvey et al., 2015) (r = 0.65, Supplementary Fig. 1c).

**Figure 1.**
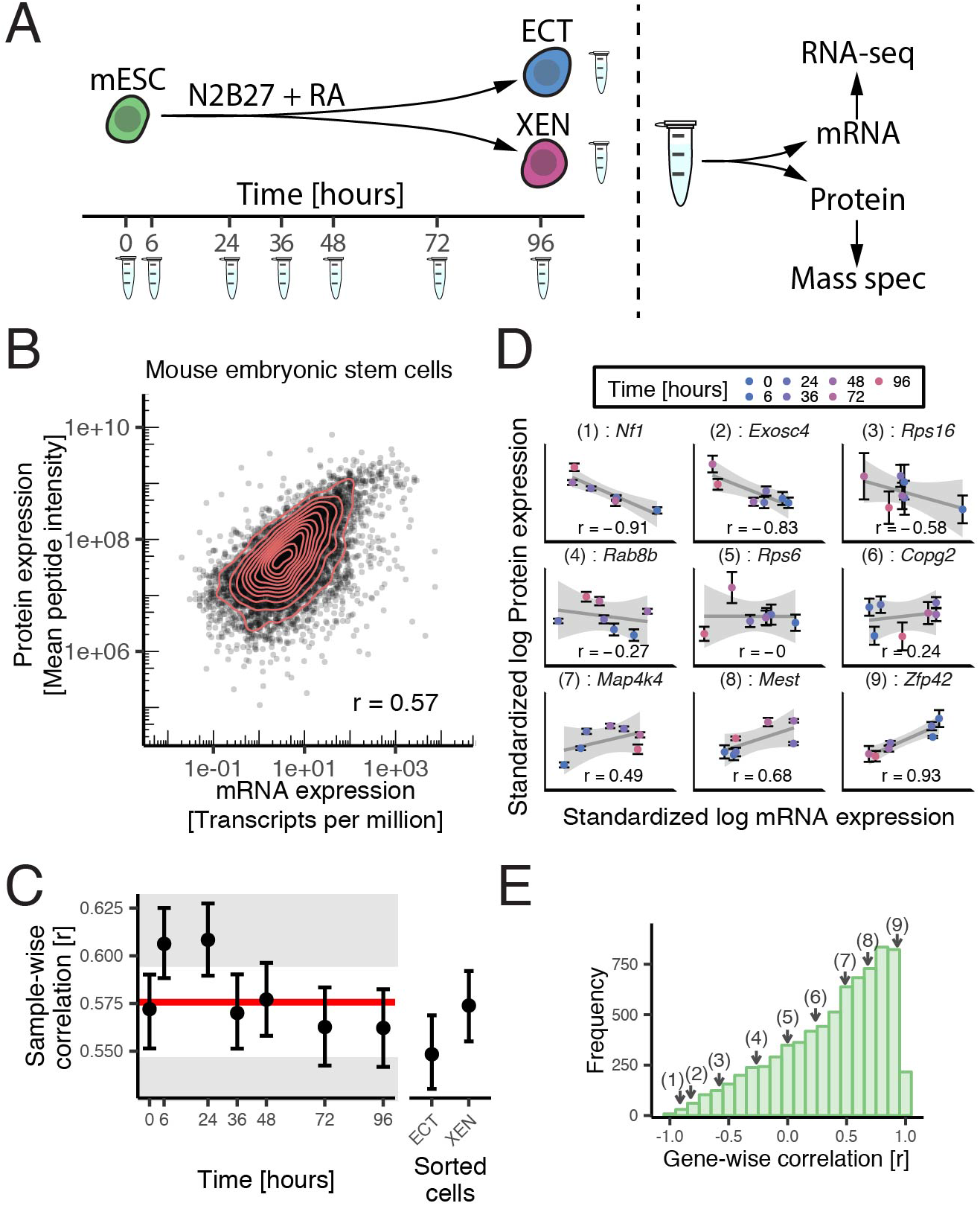
mRNA and protein expression correlate poorly during mESC differentiation. (A) Experimental setup. (B) mRNA versus protein expression of 7459 genes in mESCs. Each data point is an individual gene. Red lines indicate contour lines of equal density. (C) Sample-wise Pearson correlation between mRNA and protein for all samples. The solid line indicates the average of all time course samples. The grey area indicates the 5% rejection region for all samples being identical (see Methods). Error bars: SEM. (D) mRNA versus protein expression at all time points for nine example genes. Pearson’s correlation r is indicated for each gene. The line and grey area indicate the linear regression fit and 95% CI, respectively. Error bars: SEM. (E) Distribution of the gene-wise Pearson correlation between mRNA and protein. Numbered arrows indicate the position of the examples shown in D. See also Supplementary Figure 1.

### Correlation between mRNA and protein levels is moderate

To explore the relationship between mRNA and protein levels we first correlated the two expression levels across genes for individual time points or cell types (sample-wise correlation). In mESCs (0h time point) Pearson correlation between mRNA and protein was 0.57 (Fig. 1 b). Similar values have been reported in other mammalian systems (de Sousa Abreu et al., 2009; Jovanovic et al., 2015; Schwanhäusser et al., 2011). Sample-wise correlation was approximately the same for all samples, including the purified differentiated cell types (Fig. 1c). Low mRNA-protein correlation was thus not cell state dependent. Importantly, a low sample-wise correlation does not exclude the possibility that relative changes in protein levels during differentiation closely follow relative changes in mRNA levels. To quantify the concordance between mRNA and protein dynamics we calculated their correlation across time for individual genes (gene-wise correlation, Fig. 1d-e). Some genes, like the pluripotency factor *Rex1* (*Zfp42*) indeed exhibited a high correlation between mRNA and protein (r = 0.93 for *Rex1*). Numerous genes, like the ribosomal protein *Rps6*, for example, did not exhibit any strong correlation between protein or mRNA (r = 0 for *Rps6*). Strikingly, we also observed many genes with anti-correlated profiles, like *Arpc1a* (r = - 0.91) or *Arvcf* (r = – 0.90). Such highly negative correlations do not seem to be a result of technical noise in protein quantification, since multiple distinct peptides of the same protein show similar trends (Supplementary Fig. 1d). Overall, the distribution of gene-wise correlations, while peaking close to 1, had a long tail towards -1 (Fig. 1e). This result clearly shows that mRNA dynamics are in general not a good predictor for protein dynamics during differentiation.

### Classification by dominant temporal trends visualizes widespread discordance between mRNA and protein

Having discovered that mRNA and protein dynamics are in general dissimilar we wanted to reveal the main trends in expression dynamics and study how they differ between mRNA and protein. To that end we used singular value decomposition (SVD) to decompose an expression profile into a weighted sum of generic profiles, called eigengenes (Fig. 2a). In contrast to other classification methods, SVD allows us to discriminate systematically between the main trend (the dominant eigengene) and smaller, additional fluctuations (Fig. 2b). The first three eigengenes, which corresponded to monotonic, transient or oscillatory trends, explained 76% and 85% of the variance in mRNA and protein expression, respectively (Fig. 2c). mRNA eigengenes were more dynamic than protein eigengenes (Supplementary Fig. 1e), which reflects the buffering of mRNA dynamics by protein synthesis and degradation (Liu et al., 2016) (Jovanovic et al., 2015). Classification of all genes by their dominant mRNA and protein eigengenes (which reflect the main temporal trends) revealed widespread discordance (Fig. 2d). While there was a statistically significant enrichment of genes with similar dominant mRNA and protein eigengenes (p-value < 1E-5), most genes (60%) had discordant mRNA and protein dynamics.

**Figure 2.**
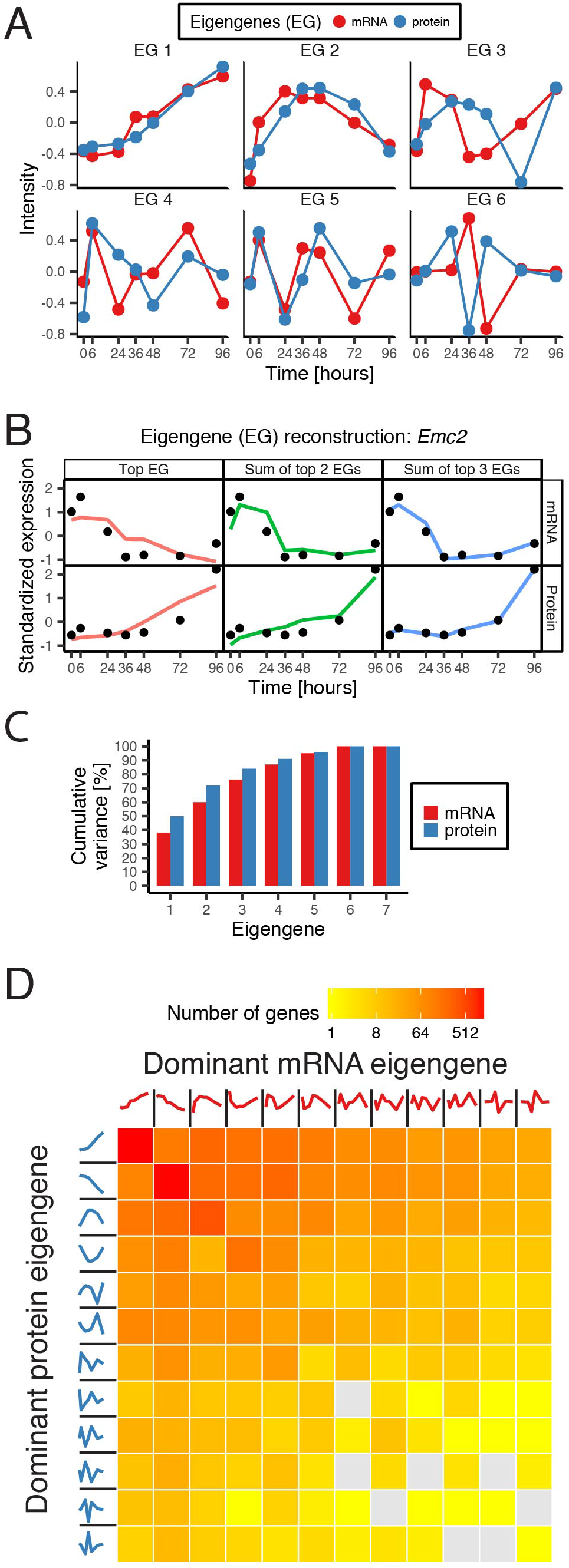
Classification of temporal mRNA and protein expression profiles by dominant trends reveals widespread discordance. (A) First six eigengenes of mRNA and protein expression profiles. (B) Reconstruction of mRNA and protein expression profiles from the top three eigengenes of an example gene. (C) Cumulative variance explained by the eigengenes for mRNA and protein profiles. (D) Classification of all genes by their dominant mRNA eigengene (columns) and protein eigengene (rows). See also Supplementary Figure 1.

### A simple kinetic model partially explains the mRNA-protein discordance for the majority of genes

The temporal delay between mRNA and protein eigengenes (Fig. 2a) sparked the hypothesis that the delay inherent to protein synthesis and degradation might cause much of the observed discordance.

To pursue this hypothesis we modeled protein turnover using a simple birth-death process with constant protein synthesis and degradation rates (Tchourine et al., 2014) (Peshkin et al., 2015) (Methods, Fig. 3a). In our model the synthesis rate k_s_ lumps all processes related to protein production (translation initiation, elongation, etc.) while the degradation rate k_d_ represents all processes leading to a reduction in protein levels (dilution due to cell division, active degradation, etc.). To avoid over-fitting, we also considered simpler models, which correspond to cases in which a protein is only synthesized, only degraded or completely constant (Fig. 3b). To select among these models, we employed the Bayesian Information Criterion (BIC), a score that penalizes the fit according to the number of parameters (Methods). To reveal whether there is a connection between a certain model and specific molecular functions, we performed GO term enrichment analysis. This analysis revealed that the “degradation only” model was enriched for genes with a role in blastocysts development and inner cell mass proliferation (Supplementary Fig. 2a). These genes are likely involved in preserving the pluripotent state, as exemplified by the pluripotency factor *Nanog*. Degradation of the corresponding proteins is crucial for the timely exit from pluripotency. GO term enrichment analysis also showed that the “synthesis only” model was enriched for genes involved in neuron development and mesenchymal cell development. These genes thus likely have specific functions in differentiated cell types and hence must be synthesized quickly to ensure proper function. An example of such a gene is *Lamb1*, which is highly expressed in XEN cells. This analysis shows that the different regulatory modes identified by our model correspond to specific functions in differentiation.

**Figure 3.**
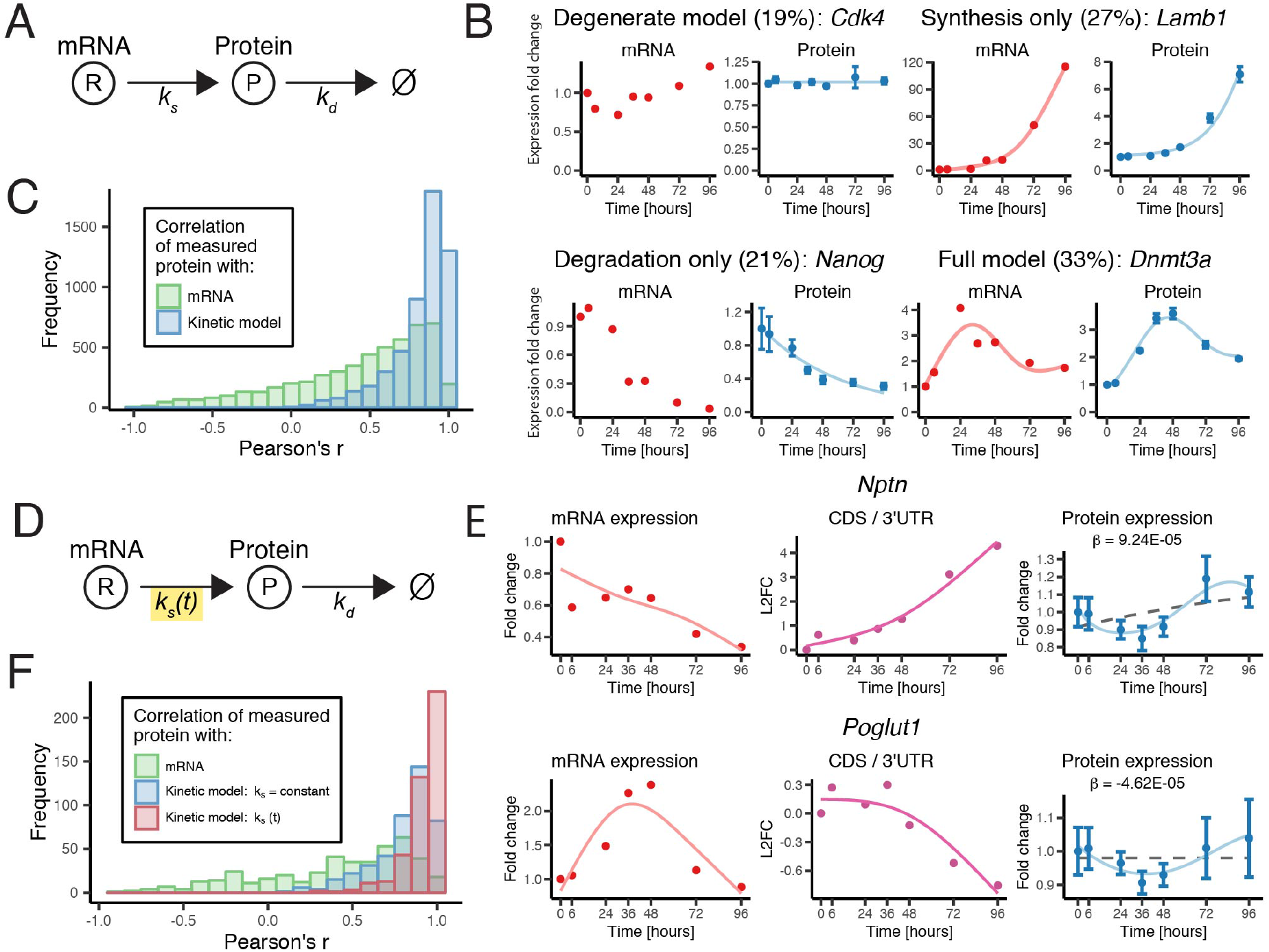
Simple kinetic models of protein synthesis and degradation explain mRNA-protein discordance. (A) Kinetic model. *k_s_* = synthesis rate constant; *k_d_*= degradation rate constant. (B) Example fits of the *full model (k_s_*>0, *k_d_* > 0) and the three reduced models: *synthesis only (k_s_ > 0, k_d_ =0), degradation only (k_s_ = 0, k_d_> 0*) and *degenerate (k_s_= k_d_=0*). Percentages indicate the fraction of genes fit best by the respective model. (C) Distribution of Pearson correlation between measured protein expression and mRNA expression or predicted protein expression. (D) Extended kinetic model. k_s_(t) = time-dependent synthesis rate. (E) mRNA expression, log ratio of expression from CDS and 3’UTR and protein expression profiles of two example genes with fits of the extended model (solid line) or the basic model (dashed line). (F) Distribution of Pearson correlation between measured protein expression and: mRNA expression, protein expression predicted by the basic model or the extended model. Error bars in (B) and (E): SEM. See also Supplementary Figures 2 and 3.

We next wanted to evaluate the validity of our model by comparison with relevant data sets from the literature. Protein half-lives (Supplementary Fig. 2b) calculated from the degradation rates were in the same range as previously reported values for other systems (Peshkin et al., 2015; Schwanhäusser et al., 2011). Synthesis rates were positively correlated with translational efficiencies determined from ribosome profiling in mESCs (Supplementary Fig. 2c) (Ingolia et al., 2011). The inferred kinetic rates are thus biologically meaningful.

In order to assess how far our kinetic model can explain the observed protein-mRNA discordance we calculated the correlation between measured and predicted protein levels (Fig. 3c). These correlations were sharply peaked close to one, which means that our simple model is able to explain a large portion of the observed mRNA-protein discordance. This discordance is likely only transient since protein-to-mRNA ratios differed most from their equilibrium value (k_eq_ = k_s_/k_d_) in the beginning but approached it over time (Supplementary Fig. 2d). This observation supports our conclusion that the observed mRNA-protein discordance during differentiation is largely a transient, dynamic imbalance caused by delayed protein synthesis and degradation.

### The CDS/ 3’UTR mRNA expression ratio is a modulator of the synthesis rate

We next sought to further refine our kinetic model and explore whether we could find predictors of protein abundance. In that respect we were intrigued by a recent report that connected the ratio of mRNA expression from the coding sequence (CDS) and 3’ untranslated region (UTR) to protein abundance (Kocabas et al., 2015). In our data sets, the CDS/3’UTR mRNA expression ratio *w* also had a non-trivial relationship with protein levels (Supplementary Fig. 2e). Consequently, we included *w* in our model as a modulator of the synthesis rate (Fig. 3d, Methods). Again, using the BIC to determine whether using an additional free parameter is warranted by the improvement of the fit, we found that 492 genes were fit optimally by the extended kinetic model (Fig. 3e). In the cases where it was optimal the extended model provided a substantial improvement over the basic model (Fig. 3f). For roughly half of those genes, *w* has a positive effect on protein synthesis and a negative effect on the other half (Supplementary Fig. 2f). While the molecular mechanism relating *w* to the protein synthesis rate is not yet known, our analysis shows that *w* is an interesting predictor that should be explored in future studies of protein dynamics.

### Failure of the kinetic model reveals dynamic post-transcriptional regulation

Despite its success in explaining the mRNA-protein discordance overall, our kinetic model does not fit the dynamics of all quantified proteins. We identified 1232 genes with a poor mRNA-protein correlation that is not appreciably improved by any of the kinetic models (Supplementary Fig. 3a). Due to the buffering of mRNA dynamics when synthesis and degradation rates are constant, the model fails in particular when the protein profile is more dynamic than the mRNA profile (Supplementary Fig. 3b). Importantly, the genes that are not fit well by our model are very similar to the full data set in their protein reliabilities (medians: 0.970 versus 0.972) and measurement errors (median SEM: 0.121 versus 0.115). Hence, technical noise is in general not the reason for the lack of a good fit. Rather, the model fails due to the assumption that kinetic rates are constant. Consequently, we consider genes that are not fit well by the model to be dynamically regulated. We sought to find sets of such genes that potentially share regulatory features. To this end we again used the classification by dominant eigengenes (Supplementary Fig. 3c). As an example, we focused on a class of genes with relatively simple dynamics: monotonically increasing mRNA and a transient increase in protein expression (highlighted in Supplementary Fig. 3c). Notably, we discovered that genes belonging to the MAPK pathway were enriched in this particular class (ConsensusPathDB, adjusted p-value = 1.8E-3, Supplementary Fig. 3d). This suggests that genes of the MAPK pathway, which is highly relevant for the differentiation of mESCs (Kunath et al., 2007), are regulated dynamically at the protein level. This analysis exemplifies that we can systematically identify sets of genes that are dynamically regulated at the protein level, likely by common mechanisms.

### Sets of genes with different functions in differentiation show distinct regulatory modes

We next wanted to concentrate further on the regulation of gene sets that are relevant for embryonic stem cell differentiation. To that end, we defined sets of markers for the pluripotent state, XEN cells, and ECT cells based on differential mRNA expression (Supplementary Fig. 4a), which were confirmed by GO term enrichment (Supplementary Fig. 4b). As a fourth gene set we considered ribosomal proteins since it has been shown previously that the translational state changes dramatically during differentiation (Sampath et al., 2008). For these 4 gene sets we calculated the average mRNA and protein profiles, correlation between mRNA and protein, classification by dominant eigengene and inferred synthesis and degradation rates for the genes that are fit optimally by the full kinetic model (Fig. 4a). This analysis of gene sets is also available on the companion website. Pluripotency markers were in general down regulated at the mRNA level (per definition) but also at the protein level. Correspondingly, we found this set to be enriched in the “degradation only” kinetic model while the “synthesis only” model is underrepresented (Supplementary Fig. 4c). This observation is consistent with the fact that pluripotency genes have to be down-regulated quickly to allow for a timely exit from pluripotency. Nevertheless, there were some genes that showed a substantial increase in protein expression and consequently had a negative correlation between measured mRNA and protein (see Supplementary Fig. 4d for examples). XEN and ECT markers were in general upregulated, where ECT markers came up before XEN markers, as shown by us previously (Semrau et al., 2016). In contrast to the set of pluripotency markers, XEN and ECT genes showed a high level of concordance between mRNA and protein, as immediately obvious from the eigengene classification. Correspondingly, both gene sets were enriched for high correlation between mRNA and protein. Additionally, XEN markers were enriched for the “synthesis only” model (Supplementary Fig. 4b). This might be related to the fact that XEN cells have to produce high levels of extracellular matrix proteins(Mulvey et al., 2015), like laminin (*Lamb1*) or collagen (*Col4a2*). Consequently, these proteins must be synthesized in a timely manner to ensure the proper function of the XEN cells. All in all, it seems that cell type specific markers defined at the mRNA level could be confirmed at the level of protein and that for these genes protein expression closely follows mRNA expression. Compared to the gene sets discussed so far, ribosomal protein (RP) genes showed a remarkable extent of discordance between mRNA and protein expression. Eigengene classification revealed that many RP genes had protein profiles that were more dynamic than their mRNA counterparts. Correspondingly, RP genes were enriched for low correlation between mRNA and protein (p-value = 3.3E-2). As cells differentiated, the protein levels of RP genes decreased, consistent with reduced cell division rates. The rate of decrease in abundance, however, was RP specific. Thus, it will be interesting to isolate ribosomes and analyze the extent to which these RP dynamics reflect ribosome remodeling and specialization (Slavov et al., 2015). In summary, we have shown that the 4 analyzed gene sets follow distinct regulatory modes that can be related to biological functions.

**Figure 4.**
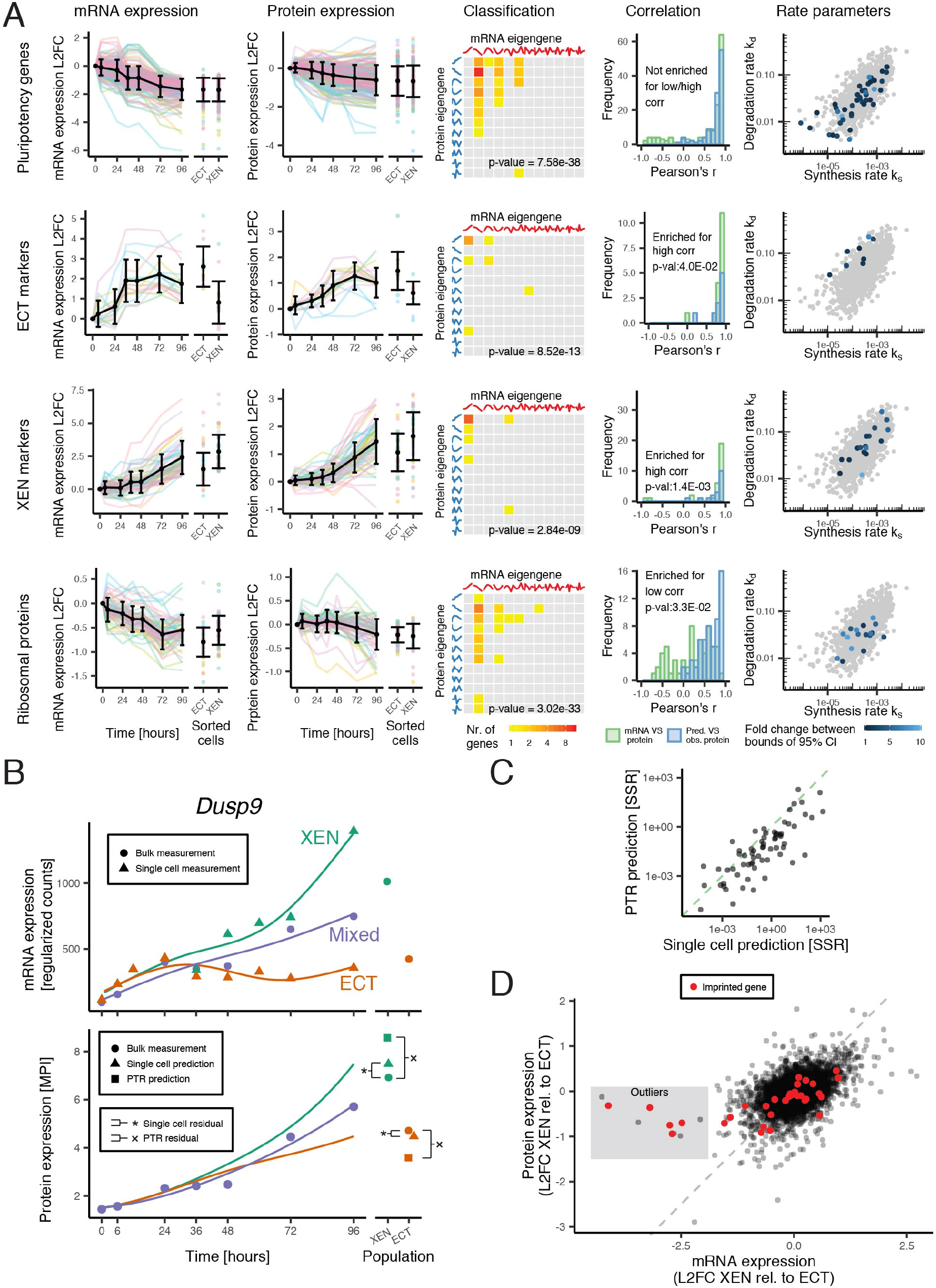
Classification and kinetic modelling reveal differences between gene sets involved in differentiation and between differentiated cell types. (A) Comparison of four gene sets that are relevant for differentiation. Log_2_ fold change (L2FC) of mRNA and protein expression are shown for individual genes (colored) and the set average (black). The p-value in the classification matrix is based on picking genes at random from all genes (chi-squared test). (B) mRNA expression of XEN and ECT subpopulations (from single cell data) and the mixed populations (bulk sample). Protein expression in XEN and ECT is predicted by applying the kinetic model to the single cell data. Alternatively, at 96 h we also predicted protein based on the protein-to-mRNA (PTR) ratio. MPI = Mean peptide intensity. (C) Sum of squared residuals (SSR) of the kinetic model-based prediction compared to the PTR-based prediction for the XEN and ECT marker genes. (D) mRNA and protein expression in XEN cells relative to ECT cells. Outlier genes are highlighted with a dark background and imprinted genes are shown in red (obtained from www.geneimprint.com, Oct-11-2016). Imprinted genes are significantly enriched in the outlier gene set (hypergeometric test: p-value = 2.72e-10). See also Supplementary Figure 4.

### The kinetic model can be applied to single-cell transcriptomics data to predict protein levels in differentiated cell types

In the experiment presented here, the existence of good antibodies for highly expressed surface markers allowed us to purify differentiated cells at 96 h and profile their proteome. For earlier time points or many other differentiation assays such an approach is difficult or even impossible. By contrast, single-cell transcriptomics methods can be applied to any differentiation system. Hence, we would like to use such data sets to predict protein levels in subpopulations. To that end, we extracted cell type specific mRNA dynamics during differentiation from our earlier single-cell RNA-seq measurement of the system (Semrau et al., 2016). We then applied our kinetic model to this data set to predict protein levels in the differentiated cell types at 96 h (Fig. 4b, Methods). Our prediction was clearly superior to a prediction that used only bulk RNA-seq measurements and protein-to-mRNA ratios (Edfors et al., 2016) (Fig. 4c). We have thus demonstrated that our kinetic model with parameters learned from bulk measurements can be applied to single-cell transcriptomics data to predict cell type specific protein levels.

We finally compared the differentiated cell types directly with each other. Overall, the correlation between mRNA and protein changes was poor and we identified a few outlier genes in particular that showed extreme behavior (Fig. 4d). These outliers had comparable protein expression in XEN and ECT cells (at most 2-fold difference) but mRNA expression was much lower in XEN cells (up to 19fold). Notably, these outliers are strongly enriched for imprinted genes (hypergeometric test, p-value = 2.3E-10). It is a well-known fact that some imprinted genes are mono-allelically expressed in extra-embryonic tissues (Miri and Varmuza, 2009). Yet, the observed down-regulation goes well beyond a two-fold change expected for mono-allelic expression. This observation demonstrates that our data set can be used to discover significant differences in gene regulation between differentiated cell types.

## Discussion

Here we systematically analyzed the dynamics of mRNA and protein expression during mESC differentiation. We observed that absolute levels of protein and mRNA are only moderately correlated in the steady (pluripotent) state, consistent with results in other mammalian systems (Schwanhäusser et al., 2011) (Wilhelm et al., 2014) (Edfors et al., 2016). Importantly, low correlation does not immediately imply a significant role of gene-specific regulation as technical noise tends to reduce the observed correlation and conventional correction schemes typically ignore the effect of systematic, correlated errors (Csárdi et al., 2015). Edfors et al. showed recently that the protein-to-mRNA ratio (PTR) for a specific gene is constant across several tissues (Edfors et al., 2016). While the PTR might allow the prediction of absolute protein levels, it is unable to capture relative changes over time or relative differences between tissues (Franks et al., 2017; Silva and Vogel, 2016).

In this study we found widespread discordance between mRNA and protein dynamics during mESCs differentiation. Such discordance has been observed recently in several systems, in particular: *Xenopus* development (Peshkin et al., 2015), *C. elegans* development (Grün et al., 2014), macrophage differentiation (Kristensen et al., 2013) and mESC differentiation (Lu et al., 2009). While this discordance is typically interpreted as a sign of (post) translational regulation (Grün et al., 2014) (Lu et al., 2009), theoretical work showed that a simple delay between mRNA and protein production can lead to a reduction in gene-wise correlation (Gedeon and Bokes, 2012) (Munsky and Neuert, 2015). Here we showed here that a simple model with constant kinetic rates, substantially reduces the discordance for 63% of discordant genes (Supplementary Fig. 3a). The same kinetic model explained protein dynamics of a third of all genes during stress response in yeast (Tchourine et al., 2014) and of 75% of all genes in *Xenopus* development (Peshkin et al., 2015). Consistently, this simple model thus explains discordance for significant proportions of the genome. We also found that the dynamics of 48% of all genes are best fit by a model that either includes only protein synthesis or degradation. A similar observation was made analyzing the stress response in yeast (Tchourine et al., 2014). We speculate that the different reduced models correspond to different regulatory mechanisms, as suggested by the enrichment of different GO terms and gene sets reported here. We further showed that protein-mRNA ratios were transiently out-of-steady-state on the way to a new equilibrium in the differentiated cell types. The observed discordance between mRNA and protein thus most likely reflects a transient, dynamic imbalance due to delayed protein synthesis and degradation. We further extended the basic kinetic model by adding the CDS-3’UTR mRNA expression ratio as a useful new predictor for the protein synthesis rate. We speculate that the underlying molecular mechanism is related to a change in the abundances of mRNA isoforms, which are believed to have different translation rates (Wong et al., 2016). Genes that were not fit well by the kinetic model, are by our definition dynamically regulated at the protein level, as constant synthesis and degradation rates are insufficient to describe the observed kinetics. This approach is complementary to the recently developed PECA method that can be used to reveal regulatory events at the mRNA and protein level (Cheng et al., 2016).

Our in-depth analysis of several gene sets revealed that cell type specific genes show a high concordance between mRNA and protein dynamics, while for RP genes the correlation is much lower. This result is reminiscent of a recent report that studied the stimulation of dendritic cells (Jovanovic et al., 2015). Jovanovic et al. found that mRNA levels explain 90% of protein fold changes after stimulation and proteins involved in the induced immune response were particularly enriched for this regulatory mode. The dynamics of “housekeeping proteins” (including RPs), on the other hand, were dominated by changes in protein synthesis and degradation rates. Similarly, Kristensen et al. reported that mRNA abundance was the best predictor for proteins that were upregulated during differentiation of monocytes to macrophage-like cells (Kristensen et al., 2013). Together with these previous reports our study supports a model in which mRNA fold changes set the level of newly produced proteins that have crucial, specific functions in the new cell state or cell type. Regulation at the level of protein turnover, on the other hand, is used to adapt the existing proteome. Importantly, we also showed that some pluripotency genes, defined as such by being down-regulated at the mRNA level, showed increasing protein expression. This result cautions against defining markers for cell states or cell types solely based on mRNA expression.

Finally, we applied our kinetic model, with model parameters learned in this study, to our earlier singlecell transcriptomics measurement of RA differentiation. Our model successfully predicted the proteomes of differentiated cell types that arise during RA differentiation. This approach thus makes it possible to measure the proteomes of cell types that cannot be purified, for example due to the lack of suitable antibodies.

In summary, this study provided the first in-depth, integrated analysis of mRNA and protein dynamics during mESC differentiation. All measured data are provided in a convenient web application. We hope that this application will facilitate future studies of specific gene sets or global relationships, for example between sequence features and protein regulation (Vogel et al., 2010).

## Author contributions

Conceptualization, S.S. and N.S.; Investigation, P. vd B., S.S., B.B. and N.S.; Resources, B.B.; Formal analysis, P. vd B. and N.S.; Software, P. vd B.; Data curation, P. vd B. and N.S.; Writing - original draft, S.S. and P. vd B.; Writing – review and editing, P. vd B., N.S. and S.S.; Supervision, S.S. and N.S.

## Acknowledgements

P. vd B. and S.S. were supported by the Netherlands Organisation for Scientific Research (NWO/OCW), as part of the Frontiers of Nanoscience (NanoFront) program. Data analysis was carried out on the Dutch national e-infrastructure with the support of SURF Foundation. N.S. was supported by a New Innovator Award from the NIGMS of the NIH under Award number DP2GM123497. We would like to thank Rudolf Jaenisch for reagents and valuable advice.

The authors declare no competing financial interests.

## Methods

### Cell culture

E14 mouse embryonic stem cells were cultured as previously described (Semrau et al., 2016). Briefly, cells were grown in modified 2i medium (Ying et al., 2008): DMEM/F12 (Life technologies) supplemented with 0.5x N2 supplement, 0.5x B27 supplement, 4mM L-glutamine (Gibco), 20 μg/ml human insulin (Sigma-Aldrich), 1x 100U/ml penicillin/streptomycin (Gibco), 1x MEM Non-Essential Amino Acids (Gibco), 7 μl 2-Mercaptoethanol (Sigma-Aldrich), 1 μM MEK inhibitor (PD0325901,Stemgent), 3 μM GSK3 inhibitor (CHIR99021, Stemgent), 1000 U/ml mouse LIF (ESGRO). Cells were passaged every other day with Accutase (Life technologies) and replated on gelatin coated tissue culture plates (Cellstar, Greiner bio-one).

### Differentiation and sample collection

Retinoic acid induced differentiation was carried out exactly as describe before (Semrau et al., 2016). Prior to differentiation cells were grown in 2i medium for at least 2 passages. Cells were seeded at 2.5 × 10^5^ per 10 cm dish and grown over night (12 h). Cells were then washed twice with PBS and differentiated in basal N2B27 medium (2i medium without the inhibitors, LIF and the additional insulin) supplemented with 0.25 μM all-trans retinoic acid (RA, Sigma-Aldrich). Spent medium was exchanged with fresh medium after 48 h.

To collect samples, cells were dissociated with Accutase. RNA was extracted from half of the sample (RNeasy, Qiagen) and the purified RNA was stored at -80C until RNA-sequencing was performed. The other half of the sample was flash frozen in liquid nitrogen and stored at −80C until mass spectrometry was performed.

### Fluorescence-activated cell sorting

FACS sorting of the differentiated cell types and quantification of the cell type frequencies was carried out exactly as described previously (Semrau *et al*., 2016).

### RNA sequencing and mRNA quantification

#### Library preparation and RNA sequencing

The libraries for RNA sequencing were prepared under standard conditions using Illumina’s TruSeq stranded mRNA sample preparation kit. The libraries were sequenced using Illumina HiSeq 3000; 40 basepair long, stranded single-end reads were sequenced at an average read depth of 40 million reads per sample. The data is available through GEO.

#### Read alignment

An RSEM-reference was created using RSEM v1.2.28 (Li and Dewey, 2011) with the Illumina iGenome GRCm38 reference using the standard settings. Next, the Illumina adapter was trimmed from the reads with *cutadapt* v1.8.3 (Martin, 2011) and low quality bases with sickle v1.33 (Joshi et al., 2011). Finally the reads were aligned with RSEM v1.2.28 (Li and Dewey, 2011) and Bowtie 2 v2.2.6 (Langmead and Salzberg, 2012) using standard settings accept for *“--sampling-for-bam --fragment-length-mean 40”*. The option “--*sampling-for-bam*” was applied so each read appears in the BAM file once. This enabled the estimation of the CDS and 3’UTR counts by *summarizeOverlaps* from the package *GenomicAlignment* v1.8.4 (Lawrence et al., 2013).

#### Gene quantification

mRNA expression was quantified by several different methods depending on the application. Transcripts per million (TPM) was calculated by *RSEM* and was used when comparing between genes since it is corrected for gene length. The more variance stabilized regularized log counts (rLC) were determined by applying the *rlog* function from *DESeq2* v1.12.3 (Love et al., 2014) on rounded expected counts obtained from *RSEM*. From this regularized counts (rC) were obtained by: rC = 2^rLC^. rLC and rC are corrected for overdispersion in low-read genes and are therefore used when comparing one gene across multiple samples. CDS and 3’UTR counts were determined by splitting the gene annotation file (GTF) with the *GenomicFeatures* package v1.26.0 (Lawrence et al., 2013) into CDS and 3’UTR for every Ensembl gene ID. Next, the number of reads on the CDS and 3’UTR features from the aligned BAM files were counted with *summarizeOverlaps* with default options. “Union”, the default option for *mode*, discards reads, if they overlap with both CDS and 3’UTR. The ratio *w* (CDS / 3’UTR) was only calculated for genes with at least 10 reads for CDS and 3’UTR in every sample.

#### Differentially expressed genes

Differentially expressed genes (DEGs) were determined by *DESeq2* v1.12.3 (Love et al., 2014) on the rounded expected counts obtained from RSEM at a false discovery rate (FDR) of 10%. The gene set ‘pluripotency genes’ were DEGs that were down-regulated when comparing the samples 0h (n=2) and 96h (n=2). XEN- and ECT-marker gene sets were DEGs that were up-regulated when comparing the samples 0h (n=2) with XEN (n=1) or ECT (n=1) respectively. Additionally, XEN- and ECT-markers have at least a 2-fold difference in expression between the two cell types.

### Mass spectrometry and protein quantification

#### Sample preparation

Pelleted cells were lysed in 400 μl RIPA buffer, except for the sorted cells, which were lysed in 200 μl RIPA buffer. Volumes of cell lysate corresponding to 100 μg protein per sample were digested with trypsin using a modified FASP protocol (Wiśniewski et al., 2009). Subsequently each sample was labeled with TMT 10-plex reagent (Prod# 90061, Thermo Fisher, San Jose, CA) according to the manufacturer’s protocol. All labeled samples were combined into a set-sample.

#### Mass spectrometry

The labeled set–sample was fractionated by electrostatic repulsion-hydrophilic interaction chromatography chromatography (ERLIC) run on an HPLC 1200 Agilent system using PolyWAX LP column (200×2.1 mm, 5 μm, 30nm, PolyLC Inc, Columbia, MD) and a fraction collector (Agilent Technologies, Santa Clara, CA). Set-samples were fractionated into a total of 40 ERLIC fractions. Each ERLIC fraction was subsequently further separated by online nano-LC and submitted for tandem mass spectrometry analysis to both LTQ OrbitrapElite or Q exactive high field (HF). One third of each fraction was injected from an auto-sampler into the trapping column (75 um column ID, 5 cm length packed with 5 um beads with 20 nm pores, from Michrom Bioresources, Inc.) and washed for 15 min; the sample was eluted to analytic column with a gradient from 2 to 32 % of buffer B (0.1 % formic acid in ACN) over 180 min gradient and fed into LTQ OrbitrapElite or Q exactive HF. The instruments were set to run in TOP 20 MS/MS mode method with dynamic exclusion. After MS1 scan in Orbitrap with 60K resolving power, each ion was submitted to an HCD MS/MS with 60K resolving power and to CID MS/MS scan subsequently. All quantification data were derived from HCD spectra.

#### Protein quantification

Relative peptide levels were estimated from reporter ion intensities measured at MS2 level. Only peptides with co-isolation below 40 % were used for quantification. The intensities of all peptides belonging to a Uniprot ID were averaged to form mean peptide intensity (MPI) for every protein. When comparing different protein samples mean peptide intensities were normalized to the sample-mean to form protein expression. Standard error of the mean (SEM) was calculated for every protein as follows: 1) for every peptide the intensities were averaged across the samples, 2) the SEM was calculated from these mean-centered peptide intensities for every protein and sample.

#### Protein reliability

The protein reliability was calculated for genes with at least two peptides quantified. For each gene, the peptides were randomly split into two groups and the MPI was calculated for each group as described above. The correlation between the MPIs of the two peptide groups across the different samples is defined as the reliability of the measurement of that protein.

### Transcriptomics and proteomics integration

While transcripts were identified by Ensembl gene IDs, Uniprot IDs were used for proteins. To integrate the two, we mapped 7681 out of 8515 Uniprot IDs to Ensembl gene IDs present in the RNA-seq data using the *idmapping* file from the Uniprot website (15-Sept-2016). An additional set of Uniprot IDs were mapped to Ensembl IDs using *biomaRt* v2.28.0 (Durinck et al., 2009). Some proteins have more than one Ensembl ID mapping to it, therefore 33 Uniprot IDs were removed, Moreover, 92 Uniprot IDs mapped non-uniquely to Ensembl IDs and for these the protein intensities were reevaluated based on Ensembl IDs. Finally, some genes were not considered because they were not detected in all samples. This resulted in a total of 7489 genes based on Ensembl gene IDs, for which we have matched mRNA and protein expression data in all samples. Additionally, we observed 3770 genes with at least 10 mRNA reads in every sample but no detected protein.

#### Sample-wise correlation

We tested if the sample-wise correlation is constant during the differentiation time course using a resampling approach. For each bootstrap a *pseudo-sample* was constructed consisting of every gene, but with mRNA and protein expression randomly sampled from the different time points. The correlations of 10,000 *pseudo-samples* were calculated to obtain a null distribution. Samples have significantly different correlation if it falls below or above the 0.36 and 99.64 percentiles of the null distribution respectively (α = 0.05, Bonferroni correction, grey area in Figure 1c).

#### Gene-wise correlation

To define a threshold for low gene-wise correlation we applied a shuffling approach (Tchourine et al., 2014). We determined the Pearson correlation for all possible permutations of the mRNA and protein expression for every gene. More than 95% of all Pearson correlation values obtained in this way were lower than 0.7, which we therefore set as the threshold between low and high correlation.

#### Expression profile classification

mRNA and protein expression were arranged in matrix form rows corresponding to genes and the columns corresponding to time course samples. These matrices were standardized by rows. Next, standard singular value decomposition (SVD) was performed separately for mRNA and protein (Wall et al., 2003). From this analysis, we obtain n eigengenes 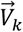 where *k* ∈,…, *n* and *n* is the number of time points. Using these eigengenes we can reconstruct the standardized expression of gene *i*, as follows: 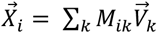, where *M_ik_* is the contribution of eigengene *k* to the standardized expression of gene *i*. We defined the eigengene with the biggest contribution to 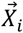 as the dominant eigengene. To determine if there is an enrichment of genes with concordant mRNA and protein eigengenes, we calculated an empirical p-value based on a null distribution generated by bootstrapped (number of bootstraps = 100,000). This null distribution was constructed under the assumption that the marginal eigengene distributions of mRNA and protein are independent. Moreover, we defined a confident set of genes with a bigger than median fold-change between the contribution of the dominant eigengene and the second most contributing eigengene for both mRNA and protein.

### Kinetic models of protein synthesis and degradation

#### Approximation of mRNA and CDS/3’UTR expression by natural cubic splines

To describe the mRNA, CDS and 3’UTR behavior in the kinetic model of protein synthesis and degradation we approximated the expression with natural cubic splines. These splines were fit on the mRNA expression and on the log_2_ fold change (L2FC) of w, which we call *ω*. The number of degrees of freedom *p* used for the fits of every gene was 4 for mRNA expression and 3 for *ω* expression. These values were automatically determined as described by Storey *et al*. (Storey, 2005). Briefly, an SVD was performed on the expression matrices of mRNA and *ω* and the first *n* eigengenes that explain at least 60% of the variance were selected. For each of these eigengenes the optimal numbei of degrees of freedom *pi* was selected by leave one out cross validation (LOOCV) and the largest *pi* was used as the number of degrees of freedom *p* to fit the natural cubic splines for all the genes of th expression matrix. The nodes of the cubic splines were equally spaced across the time course.

#### Kinetic rate parameters estimation

We model protein turnover as a birth-death process

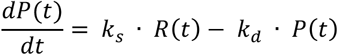

where *P*(*t*) and *R*(*t*) are protein and mRNA expression respectively. The solution of this ordinary differential equation (ODE) is given by:

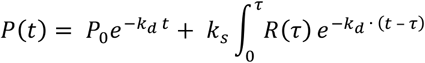

where *P*_0_ is the protein expression at t = 0 hours. The integral of this equation was estimated numerically in R using the spline fits described above. We fit the model using gene specific parameters *P*_0_, *k_s_* and *k_d_* with the Levenberg – Marquardt non-linear least squares algorithm, which is implemented in the R package *minpack.lm* v1.2-0. Additionally, we fit models where we set *k_d_* = 0, *k_s_* = 0 or *k_d_* = *k_s_* = 0. For each successful fit we determined the Bayesian Information Criterion:

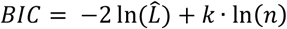

where 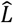 is the posterior likelihood of the fit, *k* is number of parameters in the model and *n* is the number of time points. 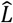 is determined by:

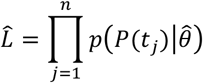

where 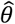 is the vector of inferred model parameters. The probabilities are estimated by assuming a normal distribution around the observed protein expression with a standard deviation equal to the SEM of the peptide intensities. The kinetic model with the lowest BIC was selected as the optimal model.

Additionally, for the subset of genes for which we could determine *ω* we constructed a model with a time-dependent synthesis rate:

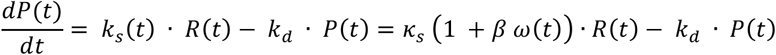

where *κ_s_* describes the constant synthesis rate and *β* parameterizes the time-dependent modulation of the synthesis rate by *ω*. The solution of this ODE:

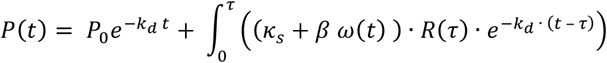

was fit to the data in the same manner as above.

#### 95% confidence region estimation

To estimate the 95% confidence intervals (CIs) for k_s_ and k_d_ we applied Wilk’s theorem:

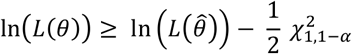

where *α* is 0.05 and 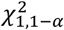 is the value at which the cumulative chi-squared distribution with 1 degree of freedom reaches 0.95. We varied k_s_ and k_d_ around the obtained fit 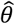 to find the edges where Wilk’s theorem holds. These edges where determined at 24 directions in the k_s_ - k_d_ solution plane to obtain a crude 95% confidence region. The projection of this region on k_s_ and k_d_ defined 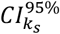 and 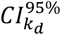, their respective 95% CIs. Note that these intervals are typically much larger than the intervals obtained when searching one parameter at a time. Genes with the full model (as determined by BIC), and with a small 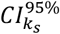 and 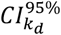 (each spanning less than a 10-fold range) were defined as the high-confidence gene set. Additionally, for genes in this set we determined the protein half-life *τ_p_* as

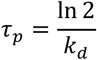

#### Protein prediction of sorted populations

We applied our kinetic model to single-cell transcriptomics data of RA driven differentiation, which we obtained previously (Semrau et al., 2016). We determined the mean expression of all cells, as well as XEN and ECT subpopulations starting from the lineage bifurcation at 36 h. All three datasets thus have identical expression up to 36 h. We then scaled the subpopulation data to the bulk data measured here for every gene in the following way: 1) We standardized the single cell time course data using the mean and standard deviation of the pooled single cell data, and 2) we scaled the standardized single cell data to the bulk data using the mean and standard deviations of the bulk time course. Next, we fit a natural cubic spline to the single cell data as before and applied the kinetic model using P_0_, k_s_ and k_d_ learned from the bulk mRNA and protein measurements. We evaluated the model performance by calculating the residuals between the predicted XEN and ECT protein expression at 96 h and the bulk measurements of protein in the purified cell types.

An alternative way of predicting protein expression is by simply multiplying a gene’s protein-to-mRNA ratio (PTR) with the gene’s mRNA expression. We defined the PTR as the mean protein expression divided by the mean mRNA expression during the time course. We predicted the protein expression of the XEN and ECT populations at 96 h using the bulk mRNA of the respective sorted populations. We used the sorted bulk data rather than the single cell data, because it is more accurate and we therefore expect this to perform better. Like with the single cell predictions, we evaluated model performance using the residuals of the PTR-predictions relative to the measured protein expression of the sorted bulk data.

### Ribosomal protein gene list

The list of RPs was compiled as all Swiss-Prot proteins curated as ribosomal proteins in their descriptions.

### Eigengene dynamics

We quantified the dynamics of the eigengene profiles as the mean of the squared second derivatives (roughness). The second derivatives were estimated numerically from three unequally spaced points by this formula:

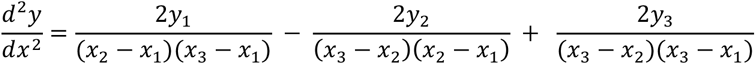

where *x*_1_, *x*_2_ and *x*_3_ are adjacent time points and *y*_O_, *y*_2_ and *y*_3_ are the respective eigengene intensities.

### GO term enrichment

GO term enrichment was performed with the R package *topGO* v2.24.0 (Alexa et al., 2006) with the *classic* algorithm. The genes were ranked using Fisher’s exact test and deemed significant with an FDR of 10%.

### Accession numbers

The RNA-seq data has been deposited in GEO (ID: GSE9563). The raw MS data has been deposited in MassIVE (ID: MSV000080461). A web application complementing this publication, which allows convenient access to all data can be found here: https://home.physics.leidenuniv.nl/~semrau/proteomics/

user name: upon request

password: upon request

## Supplementary Figures

**Supplementary Figure 1.**
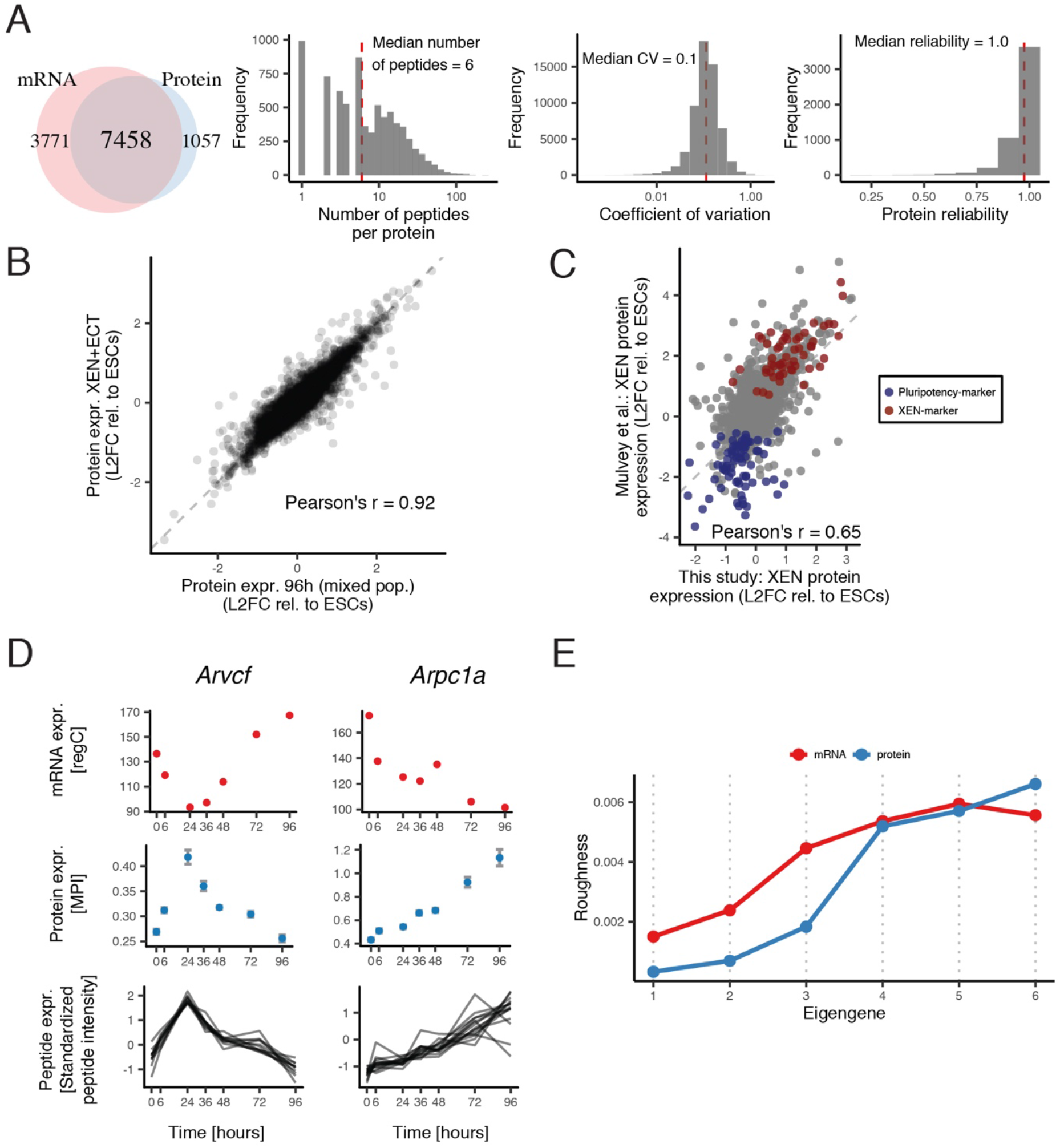
Related to figures and 1 and 2. Protein quantification using TMT labeling is robust and reproduces previous results on embryo-derived XEN cells. mRNA eigengenes are more dynamic than protein eigengenes. (A) From left to right: Venn diagram of the number of genes with quantified mRNA and protein levels (see Methods), distribution of the number of peptides used to quantify protein expression, distribution of the coefficient of variation (CV, SD/mean) of the mean-centered peptide intensities, distribution of the gene-wise protein reliability (Franks et al., 2017). The 7459 genes in the intersection are detected in all mRNA and protein samples. (B) Protein expression of the 96 h sample (consisting of both XEN and ECT cells) compared with a sample mixed *in silico* from the independently generated purified XEN and ECT cell samples. L2FC: log_2_ fold-change. (C) Protein expression in XEN cells relative to ESCs as measured in this study compared with *in vivo* derived XEN cells measured by Mulvey et al. (2015). Pluripotency- and XEN-marker gene sets were defined using a support vector machine learning algorithm. The pluripotency set is significantly enriched in genes that are downregulated in our data (p-value = 4.0E-4) and the XEN-marker gene set is enriched in genes that are upregulated (p-value = 1.4E-4, gene set enrichment analysis). (D) mRNA, protein and peptide expression for two genes with negative time-wise correlation: *Arcvf* (r = – 0.90) and *Arpc1a* (r = – 0.91). 15 and 11 peptides, respectively, were quantified for each gene. regC = regularized counts; MPI = mean peptide intensity. (E) Roughness of mRNA and protein expression eigengenes. The roughness of a profile is defined as the average squared second derivative.

**Supplementary Figure 2.**
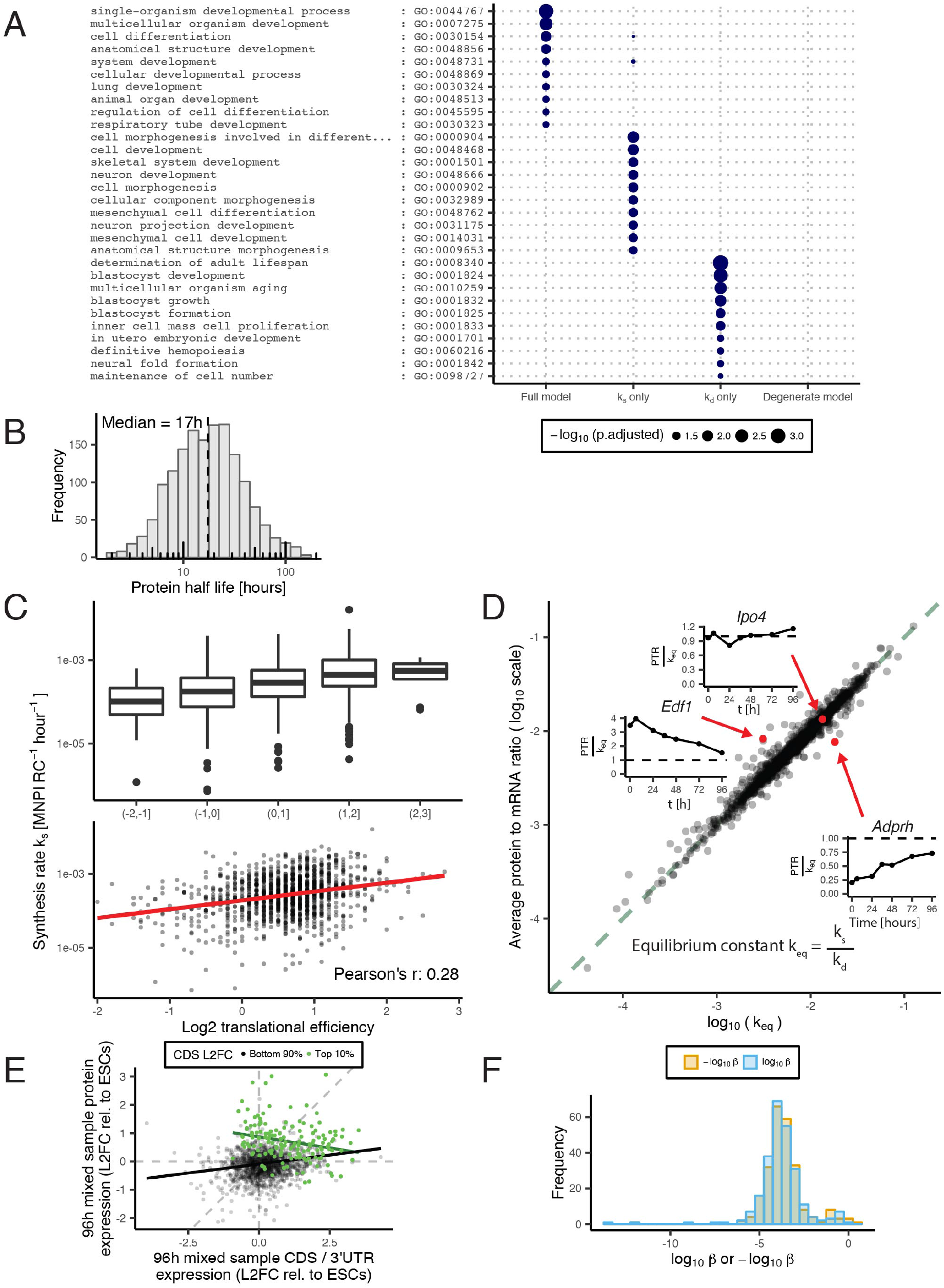
Related to figure 3. The kinetic models can be related to biological functions and the inferred kinetic rates are biologically meaningful. (A) Union of the top 10 significantly enriched *cellular differentiation* GO terms for genes fit best by each of the four kinetic models. False discovery rate = 10%. (B) Protein half life distribution for 1554 genes that were fit best by the full model (according to the BIC) and have precise estimates of the rates (upper and lower bound of the 95% confidence intervals (CIs) fall within a 10-fold range) (C) Translational efficiency (TE) in mESCs from Ingolia et al. (2011) versus our synthesis rates. We show the rates for 1284 genes (intersection between data from Ingolia et al. (2011) and the1554 genes shown in B). Boxplots represent the binned TE with whiskers indicating 1.5 x IQR. (D) Log-io protein to mRNA ratio (PTR) versus equilibrium constant (*k_eq_* =*k_s_* / *k_d_*) for the 1554 genes described in B. Each data point is an individual gene. Genes that are at equilibrium (PTR =*k_eq_*) are on the 1:1 line (green). Inserts: PTR relative to *k_eq_* across time are shown for three example genes that are above, approximately on and below the 1:1 line. (E) Ratio of CDS and 3’UTR expression versus protein expression in the 96h sample relative to ESCs. The genes with the highest CDS expression fold change are indicated in green. Solid lines indicate linear regression fits. CDS = coding DNA sequence, 3’UTR = 3’ untranslated region. (F) Distribution of the parameter β of the extended model, which sets the strength of the influence of the CDS-3’UTR ratio on the synthesis rate. Shown are the values of β for the 492 genes that are improved by the extended kinetic model (according to the BIC).

**Supplementary Figure 3.**
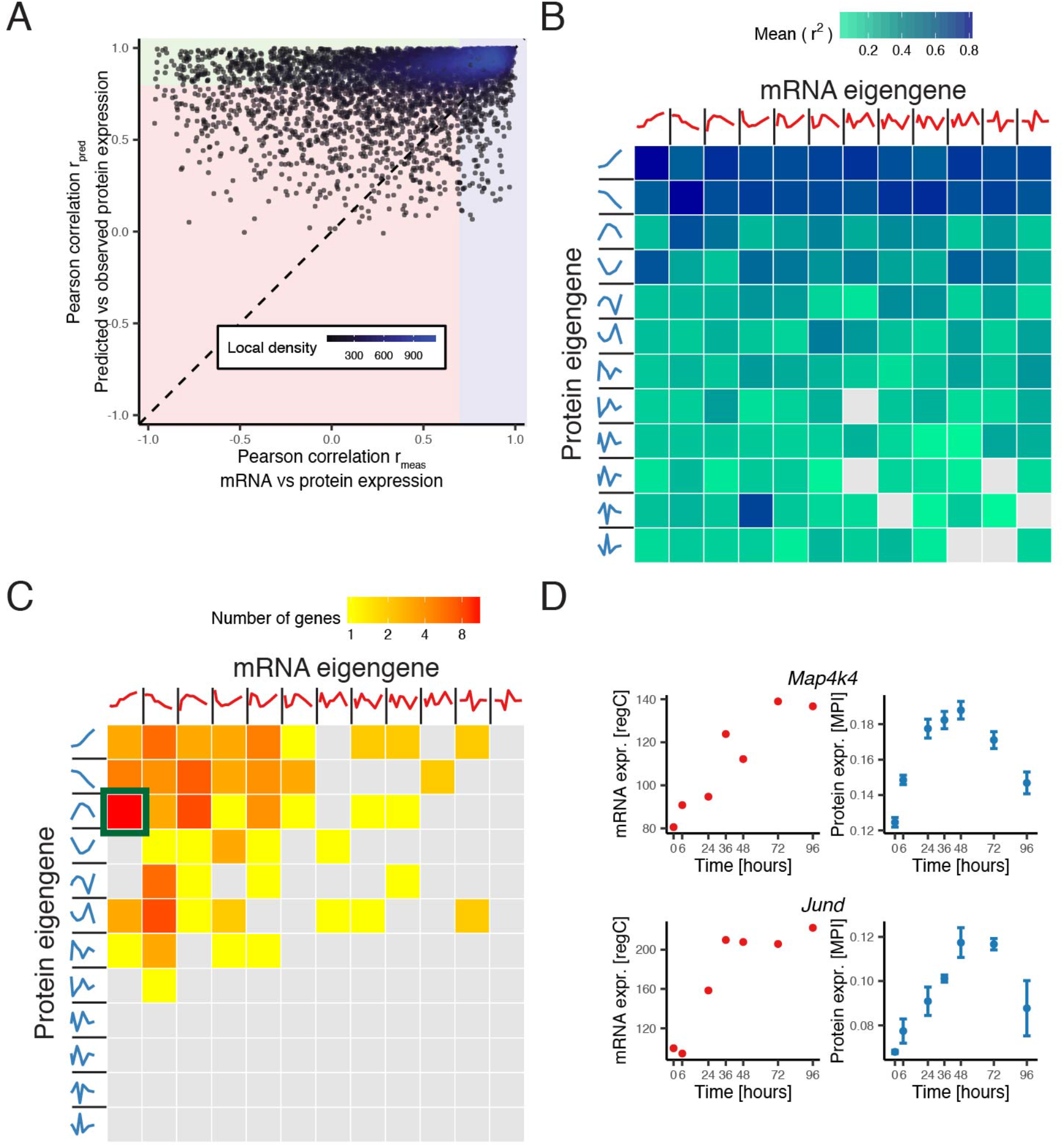
Related to figure 3. Genes in the MAPK signaling pathway are regulated dynamically at the protein level during differentiation. (A) Pearson correlation between measured protein and mRNA (r_meas_) versus Pearson correlation between measured and predicted protein (r_pred_). Background coloring indicates: concordant genes with (high r_meas_, blue), discordant genes that are not well-fit (low r_meas_, low r_pred_, red) and discordant genes that are well-fit (low r_meas_, high r_pred_, green). Here we consider genes with r_meas_ < 0.7 to be discordant (see Methods). To assure the the model prediction correlates substantially better with the measured protein than the measured mRNA we require r_pred_>= 0.8 for a gene to be considered well-fit. (B) Dominant eigengene classification of all 7459 genes. The color of a tile indicates the mean fraction of variance explained (mean r^2^) by the best-fitting kinetic model for genes with a particular combination of dominant mRNA and protein eigengene. (C) Dominant eigengene classification of the 368 genes that are not well-fit by the basic kinetic model (red area of A) and exhibit a bigger than median fold-change between the contribution of the dominant eigengene and the second most contributing eigengene. The color of a tile indicates the number of genes with a particular combination of dominant mRNA and protein eigengene. Enrichment analysis revealed an enrichment of MAPK signaling pathway genes in the tile highlighted in green (q-value = 1.8e-3). (D) mRNA and protein expression profiles of two genes from the tile highlighted in C. Error bars: SEM. regC = regularized counts; MPI = mean peptide intensity.

**Supplementary Figure 4.**
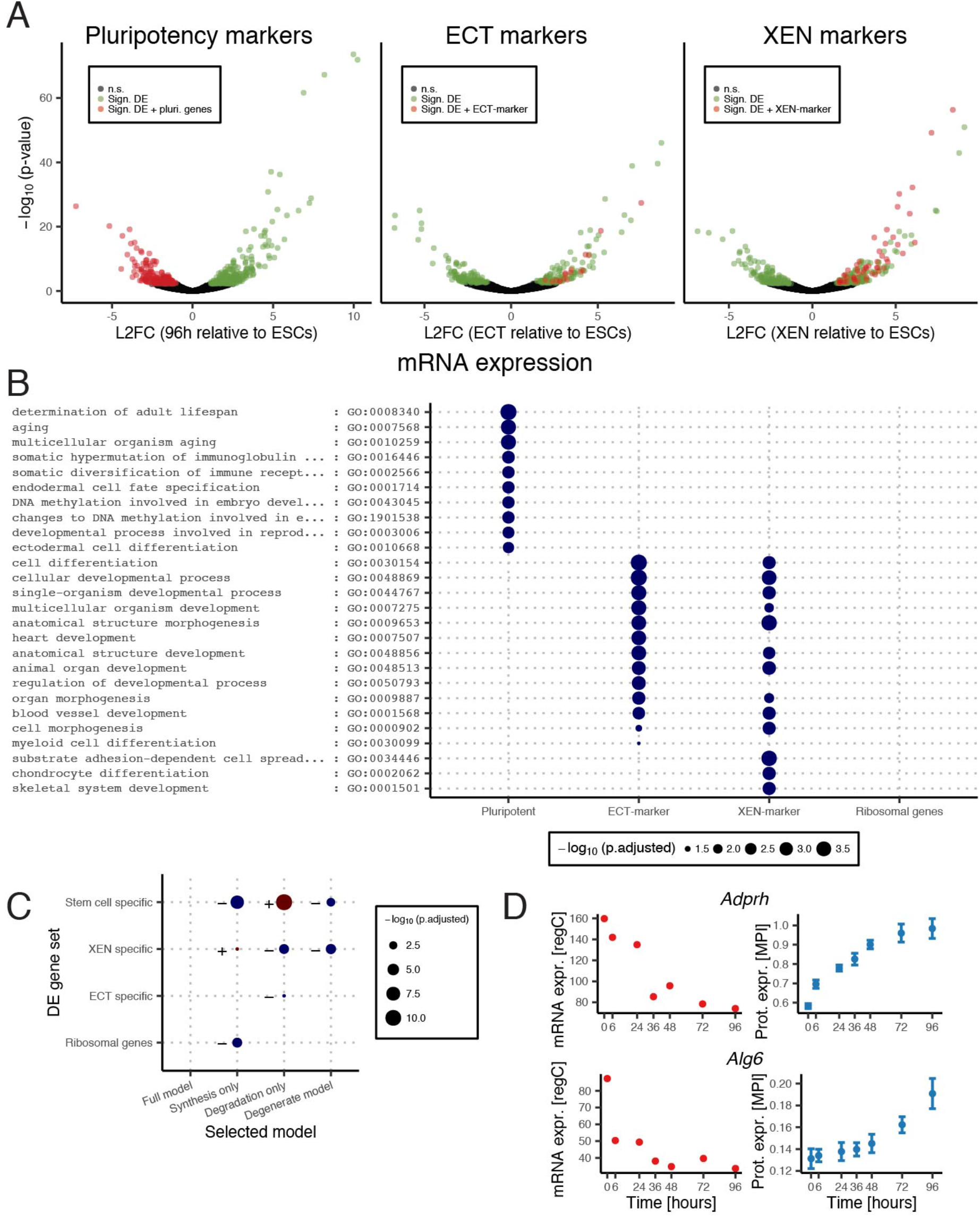
Related to figure 4. The different subtypes of the kinetic model are enriched in gene sets defined by the differentiation process. (A) Volcano plots (mRNA relative expression versus p-value for differential expression) for the 96 h sample, the ECT sample and the XEN sample. mRNA expression is always relative to the 0 h sample (ESCs). Genes colored in both red or green are significantly differentially expressed with a false discovery rate (FDR) of 10%. Only genes colored red are considered marker genes: pluripotency markers are down regulated in the 96 h sample, ECT and XEN markers are upregulated and have a minimum fold change of 2 compared with the other purified sample (see Methods). (B) Union of the top 10 significantly enriched *cellular differentiation* GO terms for genes in each of the three DE gene sets and the ribosomal genes. FDR = 10 %. (C) Overrepresentation (+ / blue) and underrepresentation (− / red) of the various subtypes of the basic kinetic model in the gene sets from B. (D) Genes in pluripotency gene set with upregulated protein expression. regC = regularized counts; MPI = mean peptide intensity.

